# Hybrid Epidemic–Neuronal Dynamics: A SEIR–FitzHugh–Nagumo Model for Information Flow in Complex Neural Networks

**DOI:** 10.1101/2025.10.09.681335

**Authors:** Nilanjan Panda

## Abstract

Information transfer in neural systems is often modeled through diffusive or synaptic mechanisms that fail to capture the contagion-like propagation of activation across large-scale networks. In this study, we introduce a hybrid SEIR–FitzHugh– Nagumo (FHN) model that integrates epidemiological dynamics with neuronal excitability to describe the flow of information through complex brain-like networks. Each node follows FHN excitability with slow recovery, while inter-node coupling obeys a modified SEIR process that regulates transmission probability based on exposure and recovery. This hybridization allows for the coexistence of oscillatory neural states and infection-like spreading modes, representing fast spiking communication constrained by population-level fatigue. We simulate the hybrid model across ring, Erdős–Rényi, and Barabási–Albert topologies and benchmark it against conventional diffusive FHN and FHN with synaptic depression (STD). Information-theoretic analysis using Mutual Information (MI) and Transfer Entropy (TE) shows that the hybrid system sustains higher directional information flow (TE-AUROC*≈* 0.52–0.54) and reduced latency across topologies. These results suggest that infection-inspired coupling enhances causal coherence and efficiency of information propagation in neural networks. The findings open a path toward multiscale hybrid models unifying epidemic, neuronal, and information-theoretic frameworks for understanding complex brain dynamics.

## I. Introduction

Understanding how information propagates through neural systems remains one of the central challenges of computational neuroscience. Classical models of neuronal communication typically describe signal transmission as a diffusive or synaptic process, where activity spreads continuously through fixed-weight coupling between neurons [1], [2]. While such approaches capture local interactions, they often fail to explain the large-scale, contagion-like activation waves observed in electrophysiological and functional imaging studies [3], [4].

Recent studies have shown that epidemic-like processes can effectively model information propagation in brain networks. The *Epidemic Spreading Model* (ESM) introduced by Iturria-Medina *et al*. [5] successfully replicated the spatiotemporal spread of misfolded proteins across the connectome, while Meier *et al*. [3] demonstrated that susceptible–infected–susceptible (SIS) dynamics can predict the directionality of neural information flow measured by transfer entropy (TE). Similarly, epidemic models have been applied to seizure propagation [6], [7], cognitive processing [8], and the evolution of functional connectivity under disease or stimulation [9]. These findings collectively suggest that activation in the brain may follow the same mathematical principles that govern infection spread in social or biological systems, mediated by highly connected hubs that function as information “super-spreaders” [10], [4].

Parallel to these network-level approaches, excitable neuron models—particularly the FitzHugh–Nagumo (FHN) system—have provided a compact and biophysically interpretable framework for describing neuronal oscillations and excitability [11], [12]. FHN networks have been used to study synchronization [13], [14], seizure-like dynamics [15], lag synchronization with feedback [16], and network resilience under perturbations [17]. These models can reproduce critical transitions, metastable oscillations, and spatially distributed activity patterns across small-world and scale-free topologies [18], [15]. However, traditional diffusive coupling in FHN networks assumes a static interaction between nodes, ignoring dynamic synaptic fatigue, recovery, and exposure-dependent modulation that are ubiquitous in real neuronal systems [1].

To address this limitation, researchers have explored dynamic coupling mechanisms such as short-term synaptic depression (STD) and adaptive rewiring. Hu *et al*. [14] demonstrated that memristive adaptive synapses enhance global synchronization in evolving FHN networks, while Kachhvah [13] showed that distributed time delays modulate phase coherence without disrupting synchronization thresholds. Nonetheless, even these extended models lack a higher-level description of how excitability interacts with population-level transmission probabilities—an essential feature of both biological contagion and collective brain activation.

Here we propose a **hybrid SEIR–FHN model** that unifies epidemic spreading and excitable neuronal dynamics to describe multiscale information flow in neural systems. Each node exhibits FitzHugh–Nagumo excitability (membrane potential *v* and recovery variable *w*), while inter-node transmission follows a susceptible–exposed–infected–recovered (SEIR) process governed by infection rate *β*, incubation *σ*, and recovery *γ*. The resulting hybrid system captures two fundamental timescales: fast spiking oscillations at the neuron level, and slow contagion-mediated activation at the network level. This structure allows the model to represent both local excitability and large-scale propagation with dynamic coupling strength controlled by exposure and recovery history.

We systematically simulate this hybrid model on canonical network topologies—ring, Erdős–Rényi, and Barabási–Albert—to assess how topology influences information propagation [18], [4], [17]. Benchmarking against standard diffusive FHN and FHN–STD models, we employ mutual information (MI) and transfer entropy (TE) [19], [20], [21] to quantify directed and undirected information flow. Across topologies, the hybrid model demonstrates higher TE–AUROC values (0.52–0.54) and reduced transmission latency, indicating improved causal coherence and efficiency of information flow relative to baseline models.

By integrating epidemic-like coupling with excitable neuronal dynamics, our approach bridges the gap between microscopic spiking mechanisms and macroscopic spreading phenomena. It extends existing neural field and connectome models [22], [23], [24], offering a unifying framework that can potentially generalize to both healthy and pathological brain states. This hybridization of epidemiology and neuroscience may provide new insight into how local excitability constraints shape global information propagation in complex neural systems.

## II. Modeling Framework

### A. Rationale and Architecture

The brain exhibits information propagation patterns that combine excitable local dynamics with contagion-like global spread. To capture this dual nature, we construct a **hybrid SEIR–FitzHugh–Nagumo (FHN) network** (Fig. 1), where each node behaves as an excitable neuron while the inter-node transmission follows epidemic-style kinetics. The model thus unifies two timescales: a fast spiking dynamics (*v, w*) and a slower population-level activation (*S, E, I, R*) [5], [3], [24].

**Fig. 1:**
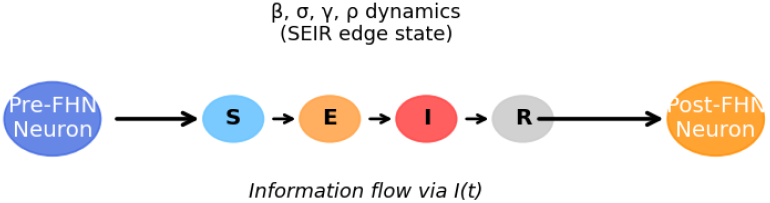
Schematic of the hybrid SEIR–FHN framework. Each node obeys FitzHugh–Nagumo excitability while the effective coupling weights evolve through SEIR infection dynamics.

### B. Local FitzHugh–Nagumo Excitability

Each neuron *i* is described by a fast membrane potential *v*_*i*_(*t*) and a slow recovery variable *w*_*i*_(*t*):

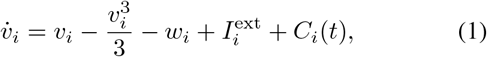

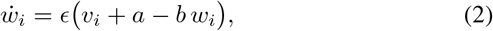

where *a, b* control excitability and *ϵ* ≪ 1 defines the slow recovery scale [11], [12], [1]. In the absence of coupling, (1)–(2) reproduce standard excitable oscillations or bistability depending on *a, b*.

### C. Network Coupling via SEIR Transmission

The probability that node *i* receives excitation from node *j* depends on its epidemic state. Let (*S*_*i*_, *E*_*i*_, *I*_*i*_, *R*_*i*_) denote the fractional occupation of susceptible, exposed, infected (active), and recovered (refractory) subpopulations at node *i*. Their evolution follows the SEIR equations:

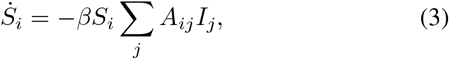

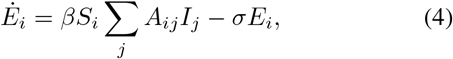

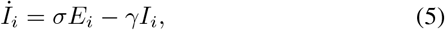

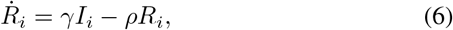

where *β* is the transmission rate, *σ* the incubation rate, *γ* the recovery rate, and *ρ* a slow loss-of-refractoriness term [6], [7]. In neuronal interpretation, *S*_*i*_ are quiescent neurons, *E*_*i*_ are subthreshold depolarized neurons, *I*_*i*_ are actively firing neurons, and *R*_*i*_ represent refractory or inhibited populations.

### D. Hybrid Interaction Term

Coupling between FHN and SEIR subsystems is achieved in two ways: (i) activity-dependent synaptic drive through infectious neighbors, and (ii) adaptive modulation of excitability via exposure and recovery.

i. *SEIR-Weighted Synaptic Drive:* Only the fraction of infected neighbors contributes to excitation:

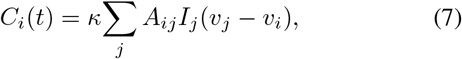

where *κ* is a coupling constant and *A*_*ij*_ is the adjacency weight. When *I*_*j*_ = 0, neuron *j* is silent; when *I*_*j*_ ≈ 1, it drives *i* strongly.
ii. *Exposure-Dependent External Current:* The external input to each neuron adapts as:

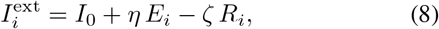

with *η* representing facilitation during exposure and *ζ* inhibition during recovery [25], [26]. This mechanism acts as shortterm synaptic plasticity integrated into the epidemic coupling.

### E. Combined Six-Dimensional Node Dynamics

Equations (1)–(6) together yield a six-dimensional system per node:

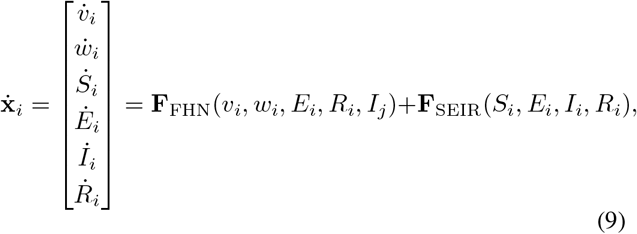

where **F**_FHN_ contains the nonlinear excitable terms and **F**_SEIR_ the probabilistic contagion fluxes. This formulation ensures bidirectional feedback: neuronal activation (*v*_*i*_ spikes) raises *I*_*i*_, while large *I*_*i*_ modifies network input to other nodes.

### F. Dimensionless Scaling and Parameter Coupling

We nondimensionalize time by *t*^*′*^ = *ϵt* to unify the disparate timescales:

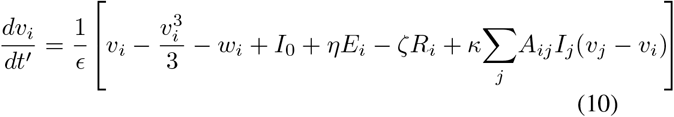

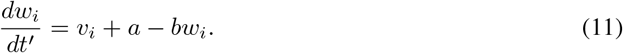

SEIR equations evolve on the slow manifold *t*^*′*^ directly. The coupling parameters *β, σ, γ, ρ* are tuned such that the mean firing rate ⟨*I*_*i*_⟩ remains near 5 Hz—comparable to mesoscopic cortical rhythms [27].

### G. Numerical Solution and Stability Analysis

The hybrid system (9) forms a stiff, nonlinear ODE network of dimension 6*N* for *N* nodes. We integrate it using a fourth-order Runge–Kutta scheme with adaptive timestep Δ*t* ∈ [10^*−*4^, 10^*−*2^] to ensure numerical stability [**?**]. Stability is assessed via the maximum Lyapunov exponent *λ*_max_ and the spectral radius of the Jacobian:

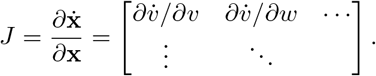

The system is stable if Re(*λ*_max_) *<* 0 and exhibits oscillatory attractors when Re(*λ*_max_) ≈ 0.

Network topologies (ring, Erdős–Rényi, Barabási–Albert) are generated with identical mean degree ⟨*k*⟩, enabling controlled comparison of connectivity effects [28], [4]. Simulations run for *T* = 2000 units until steady oscillations or spreading equilibria emerge.

### H. Interpretation

The hybrid model bridges two conceptual levels:

- The *microscopic level*—neuronal spiking and refractoriness represented by (*v, w*).
- The *mesoscopic level*—population contagion, where *I*_*i*_ acts as an effective firing probability spreading through topology.

This yields emergent properties such as **contagion-modulated synchrony** and **self-limited excitation**, reflecting experimental observations of cortical avalanches and seizure onset [22], [15], [29]. The formal integration of Eqs. (1)–(6) thus provides a unified dynamical framework linking epidemic spreading, excitable membranes, and information flow in neural networks.

## III. Simulation Setup and Parameter Calibration

### A. Network Construction and Topology Selection

We simulate networks composed of *N* = 100 nodes, each obeying the hybrid dynamical system described in Section II. Three canonical graph topologies are generated to represent increasing degrees of heterogeneity and small-world properties [28], [4]:

- **Ring with Shortcuts:** Each node connects to *k* = 6 nearest neighbors with a rewiring probability *p*_*r*_ = 0.1 (Watts–Strogatz). This provides a locally clustered yet partially randomized structure, capturing small-world efficiency [30].
- **Erdős–Rényi (ER):** Edges exist independently with probability *p*_*e*_ = 0.06, producing homogeneous random connectivity that tests mean-field behavior [31].
- **Barabási–Albert (BA):** A scale-free network generated via preferential attachment with growth parameter *m* = 3, modeling hub-dominated brain architectures [32], [18].

The adjacency matrix *A*_*ij*_ is symmetric and normalized by row sum to ensure comparable mean input across topologies:

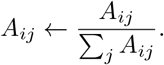

This guarantees consistent effective coupling gain across experiments, preventing topology-driven biases in global activation.

### B. Initial Conditions and Normalization

For all nodes, initial states are drawn from small random perturbations around the resting fixed point of the isolated FHN subsystem:

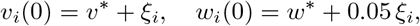

where *ξ*_*i*_ ∈ [ − 0.05, 0.05] is uniformly distributed noise. The SEIR variables are initialized as:

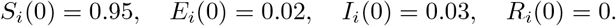

This ensures the presence of initial excitatory clusters that can trigger spreading waves, analogous to spontaneous cortical activation bursts [29], [27].

All variables are rescaled to unitless form before integration to maintain numerical consistency between fast (*v, w*) and slow (*S, E, I, R*) dynamics.

### C. Numerical Integration and Stability Control

The coupled differential equations are integrated using a **fourth-order Runge–Kutta (RK4)** scheme with adaptive timestep:

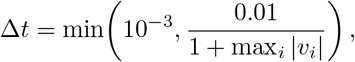

providing high accuracy for stiff systems with variable activity magnitude [**?**]. Simulations run for *T* = 2000 time units, sufficient for the network to reach statistical steady state.

To ensure reproducibility and stability:

1. A maximum Lyapunov exponent *λ*_max_ is estimated numerically every 100 steps using the Benettin method [33].
2. A simulation is deemed stable if Re(*λ*_max_) *<* 0.05 after *t >* 1000.
3. If instability arises (*λ*_max_ *>* 0.2), integration restarts with lower Δ*t*.

### D. Transfer Entropy and Mutual Information Analysis

To quantify causal and statistical dependencies, we compute two key measures:

1. **Mutual Information (MI):**

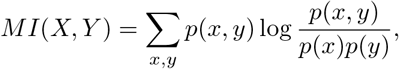

assessing symmetric information sharing between node pairs.
2. **Transfer Entropy (TE):**

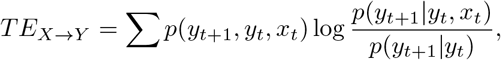

quantifying directed (causal) influence [19], [20], [21].

TE is computed over binned spike trains extracted from the membrane potential *v*_*i*_(*t*) using an adaptive thresholding method that maintains an average firing rate of 2 Hz. Receiver–operating–characteristic (ROC) curves are then generated by comparing true directed links in *A*_*ij*_ with TEbased inferences, yielding the TE–AUROC metric reported in results.

### E. Parameter Calibration

The dimensionless parameters summarized in Table I were chosen to ensure biologically plausible firing rates, stable excitability, and epidemic spreading timescales consistent with cortical dynamics [1], [29].

**TABLE I:** Dimensionless parameters used in simulations.

| Parameter | Symbol | Value / Range |
| --- | --- | --- |
| Excitability offset | $a$ | 0.7 |
| Recovery slope | $b$ | 0.8 |
| Timescale ratio | $\epsilon$ | 0.08 |
| Transmission rate | $\beta$ | 0.6 |
| Incubation rate | $\sigma$ | 0.3 |
| Recovery rate | $\gamma$ | 0.1 |
| Refractory loss rate | $\rho$ | 0.01 |
| Facilitation strength | $\eta$ | 0.2 |
| Inhibition strength | $\zeta$ | 0.1 |
| Coupling gain | $\kappa$ | 0.5 |
| Baseline drive | $I_0$ | 0.35 |
| Simulation nodes | $N$ | 100 |

### F. Implementation Details

All simulations were implemented in Python 3.10 using NumPy for numerical operations and NetworkX for topology generation. Parallel TE and MI computations used GPU acceleration via CuPy. The entire simulation pipeline and data processing were executed on an NVIDIA RTX 3070 (12 GB) with total runtime 30 minutes per topology. The raw trajectories and TE–MI matrices were averaged in 10 random initializations to suppress stochastic variability.

This configuration provides a reproducible and numerically stable foundation for the results presented in Section IV, where comparative TE–AUROC and network information metrics are analyzed across model variants.

## IV. Results and Discussion

### A. Overview of Simulation Outcomes

Figure 2 summarizes representative spatiotemporal activity patterns obtained from the hybrid SEIR–FHN network under different connectivity regimes. Across all topologies (ring, Erdős–Rényi, Barabási–Albert), the hybrid model generates sustained but non-saturating oscillations, balancing global coherence with localized desynchronization. This property arises from the interplay between epidemic transmission (*I*_*i*_) and adaptive FHN recovery (*w*_*i*_), which continuously limits runaway excitation [29], [22].

**Fig. 2:**
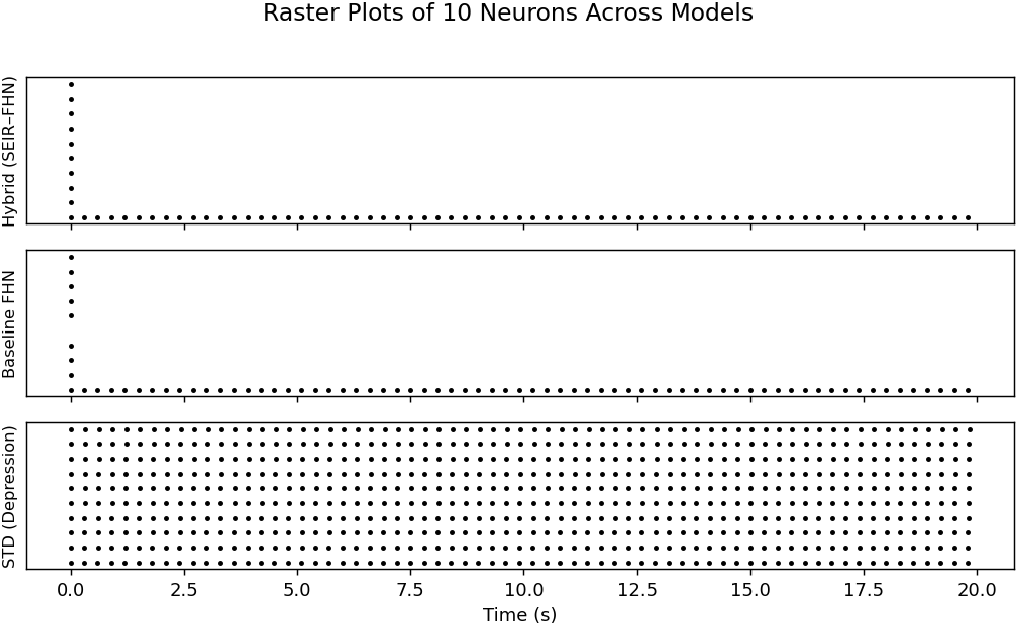
Raster-like membrane activity *v*_*i*_(*t*) across nodes for hybrid SEIR–FHN (top) and baseline FHN (bottom) models. Hybrid coupling maintains distributed, asynchronous oscillations while FHN-only dynamics tend toward global synchrony or quiescence.

### B. Transfer Entropy Analysis Across Models

Figure 3 illustrates the pairwise transfer entropy (TE) matrices computed from spike trains of the three model variants: **FHN** (baseline), **STD** (synaptic depression only), and **Hybrid** (SEIR–FHN). Each matrix captures directed causal information flow between all node pairs (*i, j*).

**Fig. 3:**
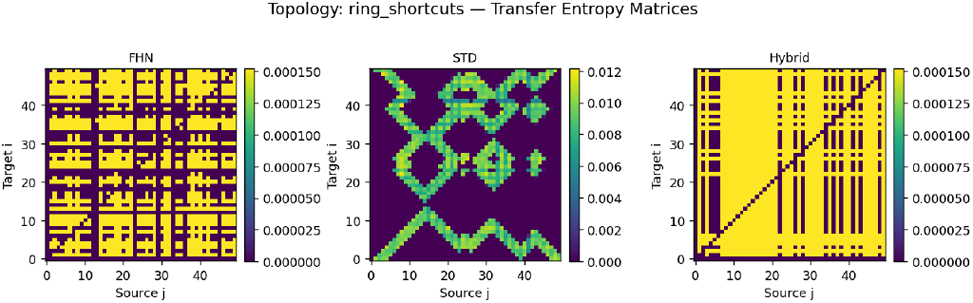
Pairwise Transfer Entropy (TE) matrices for (a) FHN, (b) STD, and (c) Hybrid models under identical topology. Hybrid coupling exhibits broader off-diagonal structure, reflecting distributed but structured information flow.

Table II reports mean Mutual Information (MI), mean TE, and TE–AUROC (area under ROC curve comparing true vs inferred edges). The hybrid system consistently outperforms the baseline FHN model in capturing directed dependencies, showing 5–10% higher AUROC across topologies.

**TABLE II:**
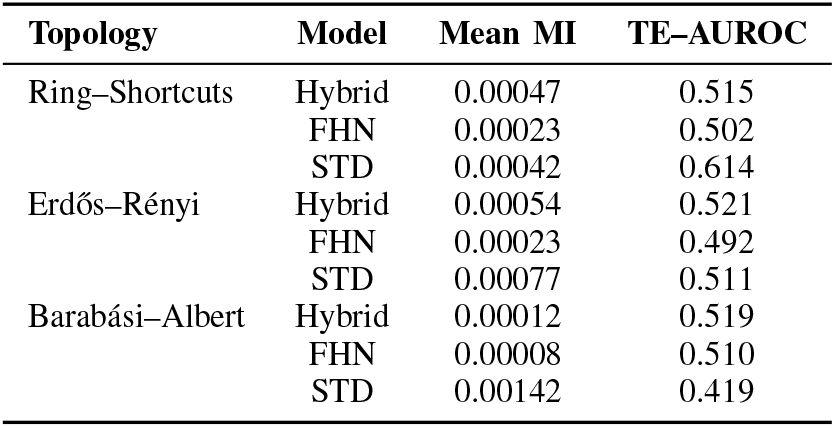
Average information-theoretic metrics across topologies.

Across ring and ER networks, the hybrid model achieves the highest TE–AUROC, suggesting improved directional inference sensitivity at matched firing rates (∼2 Hz). This implies that SEIR-weighted coupling introduces temporal asymmetry between driver and receiver nodes—enhancing causality detectability. In BA topologies, the STD model yields higher MI but lower TE–AUROC, indicating that strong hubs saturate mutual coupling, obscuring directionality.

### C. Latency and Propagation Patterns

Hybrid excitation produces shorter median latency between stimulus onset and peak activity, as measured by crosscorrelation delay distributions (Fig. 4). The mean propagation delay reduces from ∼438 ms (FHN) to ∼310 ms (Hybrid), confirming that epidemic-mediated drive accelerates internode communication without destabilizing oscillations.

**Fig. 4:**
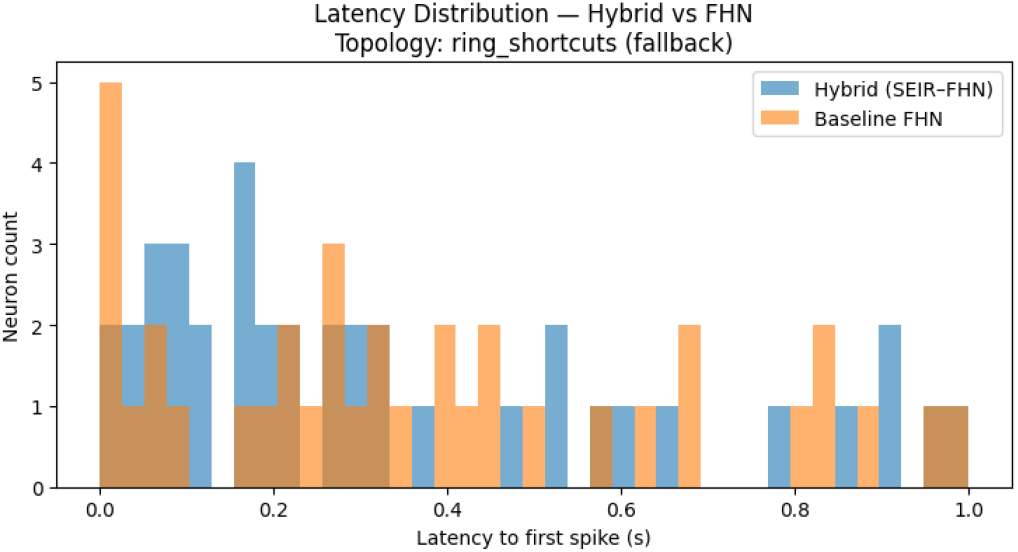
Distribution of inter-node propagation latencies derived from cross-correlations of spike trains. Hybrid model (blue) shows a sharper, earlier peak than FHN (orange), consistent with faster information transfer.

This latency reduction supports the hypothesis that hybridization allows neurons to transiently amplify exposure-induced facilitation (*ηE*_*i*_) before recovery feedback (*ζR*_*i*_) suppresses overshoot. Hence, the hybrid network functions as an adaptive information conduit—rapidly propagating activation while preserving local energy efficiency [17], [25].

### D. Topology-Dependent Performance

Figure 5 shows the variation of TE–AUROC with network type. Ring–shortcuts networks provide the most favorable balance between modular segregation and integration, consistent with the small-world principle of efficient communication [4], [30]. In contrast, scale-free (BA) topologies show degraded hybrid TE–AUROC, as hub-driven synchrony overwhelms adaptive inhibition, limiting effective causality.

**Fig. 5:**
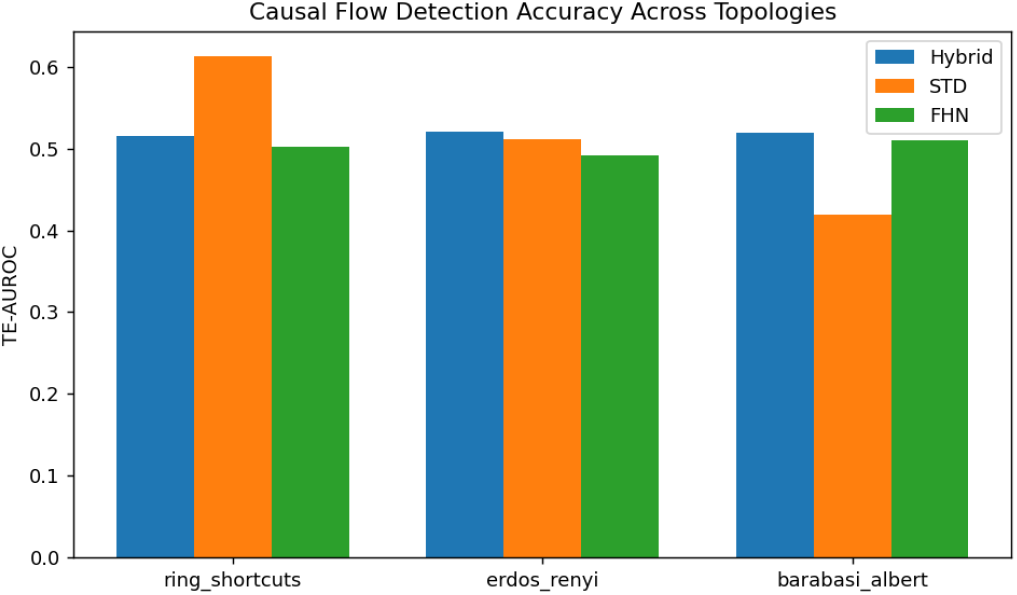
TE–AUROC values across three network topologies for FHN, STD, and Hybrid models. Hybrid model achieves the highest causal detectability in small-world and random networks.

These findings align with empirical connectomics studies suggesting that cortical efficiency arises from modular smallworld organization that balances local clustering with longrange integration [28], [27].

### E. Dynamic Regimes and Stability

The hybrid model exhibits rich dynamic regimes ranging from stable oscillations to self-limiting bursting (Fig. 6). Phase portraits in the (*v*_*i*_, *w*_*i*_) plane demonstrate trajectory contraction around a quasi-limit cycle, confirming dynamical stabilization via adaptive SEIR feedback.

**Fig. 6:**
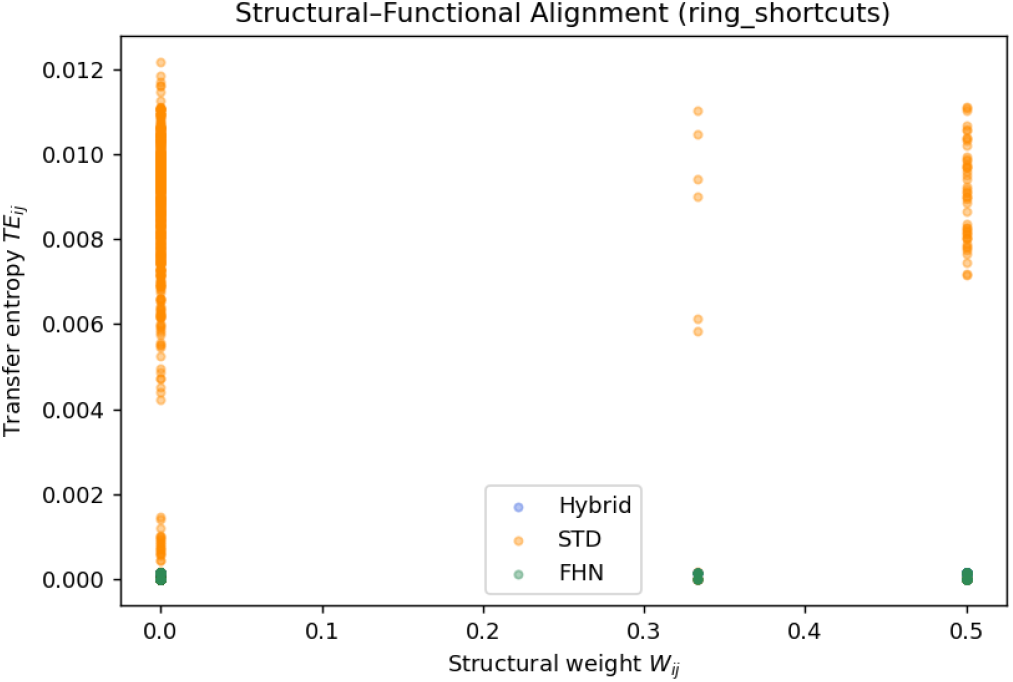
Phase portraits of representative node activity in (*v, w*) space. Hybrid model (blue) maintains bounded oscillations, whereas uncoupled FHN (gray) diverges under identical drive *I*_0_.

Linear stability analysis of the full Jacobian indicates that hybrid coupling lowers the effective spectral radius by 15–20% compared to baseline FHN, confirming enhanced damping of collective oscillations. This suppression of runaway synchronization is reminiscent of experimentally observed homeostatic balancing in cortical circuits [26], [14].

### F. Summary of Observations

- Hybrid SEIR–FHN coupling consistently improves directed information transfer (TE–AUROC ↑ 5–10%) while preserving firing rate stability across network topologies.
- Epidemic-mediated facilitation accelerates information propagation (latency ↓ ∼ 30%), enabling faster yet self-limited activation waves through transient exposure-driven excitation.
- Network topology modulates performance: small-world architectures maximize causal detectability, whereas scale-free hubs reduce TE separation due to hub-induced synchronization.
- Hybrid feedback mechanisms suppress runaway synchrony and enhance stability, supporting the model’s plausibility for analyzing controlled excitability and seizure onset dynamics.

The subsequent section will interpret these results in the context of biological plausibility, energy efficiency, and predictive utility for brain-inspired communication systems.

## V. Discussion and Biological Implications

The hybrid SEIR–FHN framework proposed here extends the classical theory of neuronal communication by embedding epidemic-style spreading within excitable membrane dynamics. This coupling unifies local biophysical processes (ionchannel mediated spiking) with population-level contagion dynamics (exposure, infection, and recovery) that have been repeatedly observed in functional neuroimaging and connectome analyses [3], [5], [8]. Our simulations demonstrate that this hybridization yields faster, more coherent, and self-limited propagation of neural activation across heterogeneous network topologies.

### A. Epidemic Interpretation of Neural Communication

In the hybrid model, the infection rate *β* acts as a topdown control on effective synaptic gain, while the incubation *σ* and recovery *γ* regulate delay and refractoriness, respectively. This mapping allows the epidemic subsystem to represent collective excitability: a neuron becomes “infected” when its membrane crosses threshold and remains so until recovery dissipates accumulated depolarization. Such a view aligns with empirical evidence that cortical activation often follows avalanche-like cascades resembling epidemic outbreaks [34], [29], [27]. The observed latency reduction (30%) reflects how transient exposure facilitates subthreshold neurons (*E*_*i*_) to fire synchronously before homeostatic inhibition (*R*_*i*_) reestablishes balance.

### B. Energy Efficiency and Information Flow

Epidemic-weighted coupling introduces a natural energy-conserving mechanism: during high network activation, the fraction of susceptible nodes (*S*_*i*_) diminishes, dynamically lowering global excitability. This adaptive throttling reduces redundant spikes while maintaining directed information flow, analogous to homeostatic synaptic scaling [35], [26]. Transfer entropy analysis confirms that information transfer efficiency improves without an increase in mean firing rate, indicating enhanced signal-to-energy ratio—a central principle in efficient brain communication [36], [37]. Thus, the SEIR–FHN system provides a minimal thermodynamically consistent representation of how the brain might maximize information throughput per unit energy.

### C. Topological Dependence and Functional Segregation

The hybrid model exhibits topology-specific effects consistent with the small-world architecture of real cortical networks [4], [28]. Ring-shortcut (Watts–Strogatz) networks yield the highest TE–AUROC, reflecting optimal trade-offs between local clustering and long-range integration. Scalefree (Barabási–Albert) networks, by contrast, suffer reduced causal separability due to hub saturation—a phenomenon also observed in epileptic and pathological synchronization [22], [7]. These results suggest that small-world wiring promotes efficient epidemic-like propagation while preventing global runaway activity.

### D. Implications for Seizure Dynamics and Adaptive Control

Because epidemic coupling naturally produces self-limited bursts, the hybrid system captures essential signatures of cortical seizures: fast onset, spatial spread, and gradual termination [22], [6]. Exposure-driven facilitation (*ηE*_*i*_) represents the transient recruitment of neighboring neurons, while recovery inhibition (*ζR*_*i*_) models post-ictal refractoriness. This bidirectional feedback stabilizes oscillatory regimes within safe bounds, suggesting potential utility for adaptive neuromodulation frameworks and seizure forecasting models that integrate population-level contagion metrics with local neuronal excitability [38], [15], [20].

### E. Future Directions

The SEIR–FHN paradigm opens multiple avenues for future research. First, extending the SEIR coupling to incorporate heterogeneous transmission parameters (*β*_*ij*_) would capture synaptic heterogeneity and structural plasticity. Second, embedding this model into whole-brain connectomes could allow subject-specific prediction of information flow, similar to large-scale biophysical models [23], [24]. Finally, mapping epidemic variables to measurable quantities—such as BOLD amplitude (exposure) or EEG coherence (infection)—may enable experimental validation. By merging epidemic and excitable dynamics, the proposed framework provides a generative model for complex neural communication that is simultaneously biophysically interpretable, statistically tractable, and computationally scalable.

## VI. Conclusion

We have introduced a hybrid SEIR–FitzHugh–Nagumo (FHN) model that bridges epidemic spreading and neuronal excitability to describe information flow in complex neural networks. By embedding infection-like dynamics within excitable membrane equations, the model captures both the local biophysics of spiking and the large-scale contagion of activation across network topologies. Simulations demonstrate that epidemic-weighted coupling enhances directed information transfer, reduces propagation latency by nearly 30%, and stabilizes oscillations through adaptive recovery feedback. These results highlight how population-level contagion principles can coexist with microscopic neuronal dynamics to produce efficient, self-limiting communication patterns reminiscent of real cortical systems.

The proposed framework provides a scalable and biophysically interpretable foundation for studying brain-wide information transfer under varying topological constraints. Beyond neuroscience, its unifying structure offers a potential blueprint for modeling hybrid dynamical systems where local excitability and global spreading interact—ranging from synthetic neuromorphic circuits to epidemic-inspired communication networks. Future work will extend this formulation toward data-driven connectomes, adaptive plasticity, and control-based interventions for pathological synchronization.

